# Seasonal IVIG contains high infectivity neutralizing and hemagglutination inhibition activity against seasonal flu virus strains selected for vaccines (2020-22)

**DOI:** 10.1101/2022.01.31.478469

**Authors:** José María Díez, Daniel Casals, Carolina Romero, Rodrigo Gajardo

## Abstract

IVIG solutions (10%) were tested for infectivity neutralization and hemagglutination inhibition against the influenza strains (two A and two B) recommended by the WHO for inclusion in vaccines for Northern and Southern hemispheres 2020-2022. The IVIG batches used were manufactured (June 2020) from plasma pools collected in the US. Potent neutralization and inhibition of hemagglutination were seen with these IVIG batches. This raises the possibility that IVIG could be added to existing therapies, especially in immunocompromised patients, to fight serious influenza infections. Further study of the potential of IVIG in influenza is warranted.

## Introduction

Seasonal influenza viruses are a major global public health problem year after year with the possible exception of this past year due to anti-COVID-19 pandemic safety measures. Recent global estimates of the mortality burden of influenza are in range of 290,000-650,000 deaths annually. ^1, 2^ The flu season in most of the world was very mild in 2020-2021. ^3^ As previously mentioned, this was likely due in part to the public health measures implemented to control the COVID-19 pandemic (e.g., masking, physical distancing, restrictions on large indoor and outdoor gatherings). ^4, 5^ With the foreseeable decline in COVID-19 cases in many parts of the world, the public health measures in place for the last two years would be lifted. This could lead to a rebound in influenza cases for the 2021-2022 season and thereafter with case levels well above those expected for a normal flu season. ^3^

Vaccination is currently one of the best available countermeasures against influenza infection. Antiviral drugs complement vaccination as both treatment and prophylaxis. Influenza infections still result in significant morbidity and mortality in immunocompromised patients. Other treatment options are needed that can be used in combination with current therapies.

Since intravenous immunoglobulin (IVIG) products are potentially a reflection of the immunological history of the donor population (they are derived from plasma of thousands of individual donors). Normal seasonal IVIG solutions were tested for antibodies against the influenza viruses selected by the World Health Organization (WHO) for inclusion in seasonal flu vaccines for 2020-2022. IVIG solutions are approved medical products manufactured from pooled plasma collected from thousands of healthy donors. Donors of different age groups and from different geographical areas may have unique histories regarding influenza infections. The antibodies in the products manufactured from plasma donated by these populations would reflect antibodies arising from both natural infections and immunizations in recent years.

In this study, IVIG solutions manufactured in June 2020 (from plasma collected in the prior 6-months) were tested for activity against the four influenza strains recommended in February 2020 for the Northern Hemisphere 2020-2021 influenza season,^6^ the strains subsequently recommended in September 2020 for the Southern Hemisphere 2021 influenza season ^7^ and in February 2021 for the Northern Hemisphere 2021-2022 season. ^8^ Activity was measured using neutralization and hemagglutination inhibition assays.

## Materials and Methods

### Influenza virus strains

The influenza viruses used in these studies were obtained from National Institute for Biological Standards and Controls NIBSC (Ridge, Herts, UK). Viral strains selected by the WHO for the quadrivalent influenza vaccine usually contain two A strains (H1N1 and H3N2) and two B strains (Victoria lineage and Yamagata lineage). The four virus strains selected by the WHO for the Northern Hemisphere 2020-2021 influenza season were: A - Guangdong-Maonan/SWL1536/2019 (H1N1)pdm09-like; A - Hong Kong/2671/2019 (H3N2)-like; B – Washington/02/2019 (Victoria lineage)-like; B - Phuket/3073/2013 (Yamagata lineage)-like. ^6^ The four virus strains recommended for the Southern Hemisphere 2021 influenza season were the same as those listed above except for the A/H1N1 strain which was Victoria/2570/2019 (H1N1)pdm09-like virus. ^7^ The four recommended virus strains for the Northern Hemisphere’s 2021-2022 influenza season were: A - Victoria/2570/2019 (H1N1)pdm09-like virus; A - Cambodia/e0826360/2020 (H3N2)-like virus; B - Washington/02/2019 (B/Victoria lineage)-like virus; and B - Phuket/3073/2013 (B/Yamagata lineage)-like virus. Three of these strains were recommended in the previous vaccines, with the novel strain being the Cambodia/e0826360/2020 (H3N2)-like virus. ^8^

### Immunoglobulin products

IVIG solutions (10%), Flebogamma DIF (batch A4GLE00311) and Gamunex (batches B2GGEOOO63 and B1GJE00153), manufactured in June 2020 by Grifols (Barcelona, Spain, and Clayton NC; USA, respectively) were used in this study.

### Microneutralization

Viral neutralization studies were performed as described in the supplementary material. The percent neutralizing activity was determined by calculating the average of the positive control wells (virus and cells only – equivalent to 0% neutralization) and subtracting that value from the average value of the negative controls (cells and culture medium only equivalent to 100% neutralization) to obtain the response range. The values from the test wells minus the negative control were expressed as a percentage of the response range (negative control – positive control).

Percent neutralization was plotted against the log dilution of IVIG using Prism software (GraphPad, San Diego, CA, USA). ID50 (the dilution producing 50 percent neutralization) and IC50 values (the concentration producing 50 percent neutralization) were calculated using the same software.

### Inhibition of hemagglutination

Assessment of hemagglutination inhibition was performed using a solution of 0.75% chicken (*Gallus gallus domesticus*) erythrocytes (Innovative Research, Novi, MI, USA; Seguridad y Salud Animal SL.,Barcelona, Spain) diluted in Alseve?s solution (Sigma-Aldrich, St. Louis, MO, USA) as previously published ^9, 10^.

One hemagglutination unit (HU) was considered the highest dilution of the virus that produces erythrocyte agglutination. After titrating the influenza virus strains, suspensions of 4 HU (in 50 μL) of the virus were added to 96-well plates (U- or V-shaped wells) except for negative control and IVIG control wells. IVIG products to be tested were diluted in phosphate-buffered saline (PBS: Life Technologies/Thermo Fisher Scientific, Waltham, MA, USA). IVIG dilutions (50 μL) was added to the test wells (eight replicates). In the negative control wells, 100 μL of PBS was added. In the IVIG control wells, 50 μL of the highest concentration and 50 μL of PBS were added.

After one hour of incubation at 37°C, the 0.75% solution of chicken erythrocytes (25 μL) was added to all wells. The plates were incubated for one hour at room temperature and then read. Results were reported as the highest dilution of IVIG that inhibited hemagglutination.

## Results

Table 1 shows the neutralization and hemagglutination inhibition titers of seasonal IVIG (June 2020) tested against the influenza strains selected by the WHO for the flu seasons in the northern and southern hemispheres in 2020-2022. These results show that the IVIG solutions produced in June 2020 contained antibodies against the four influenza strains recommended by the WHO for the northern hemisphere 2020-2021 flu season. These data also show that the same IVIG solutions (June 2020) showed similar activity against the different flu strain recommended by the WHO for the southern hemisphere 2021 season and the different strain recommended for the northern hemisphere 2021-2022 season.

**Table 1.**
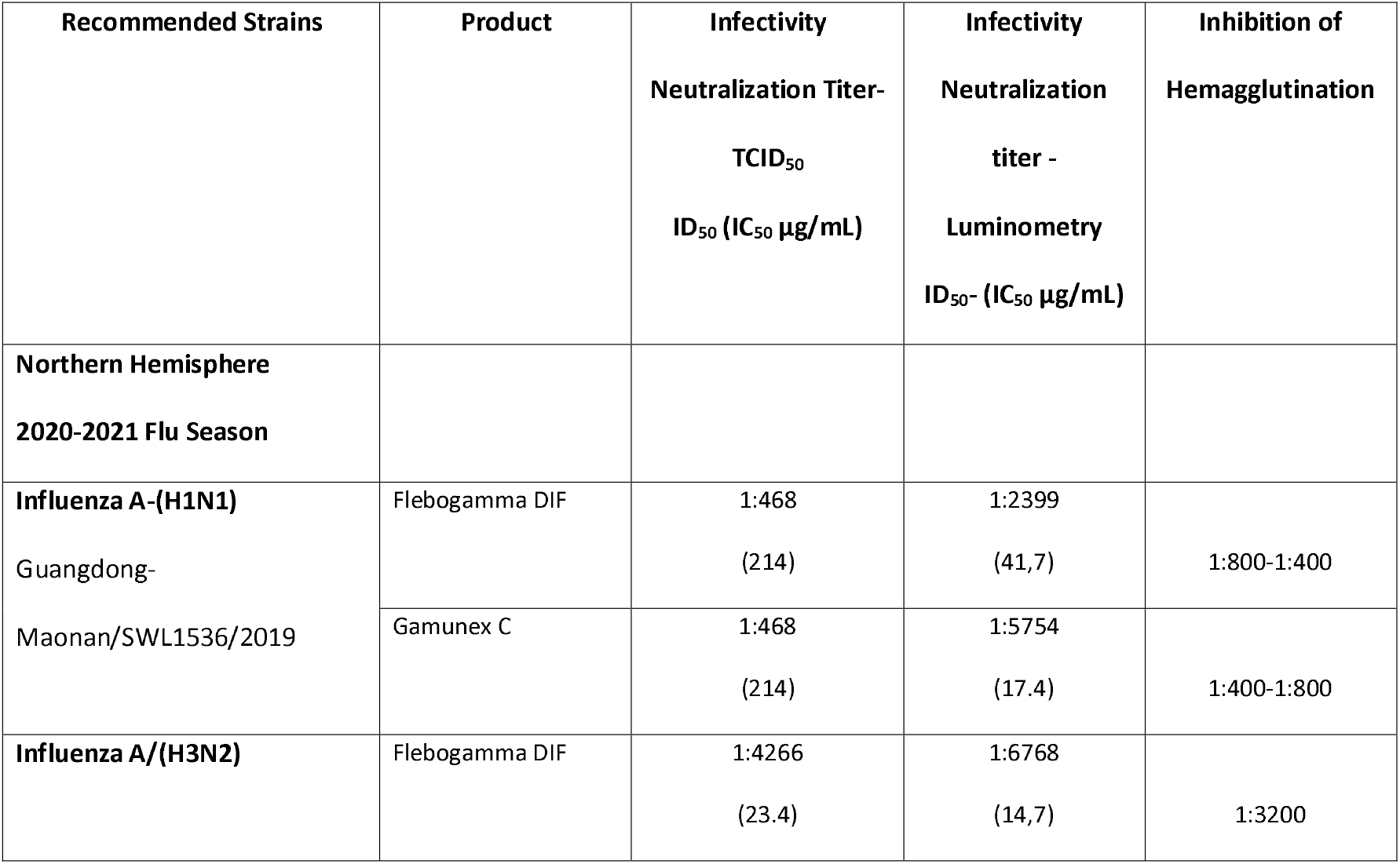
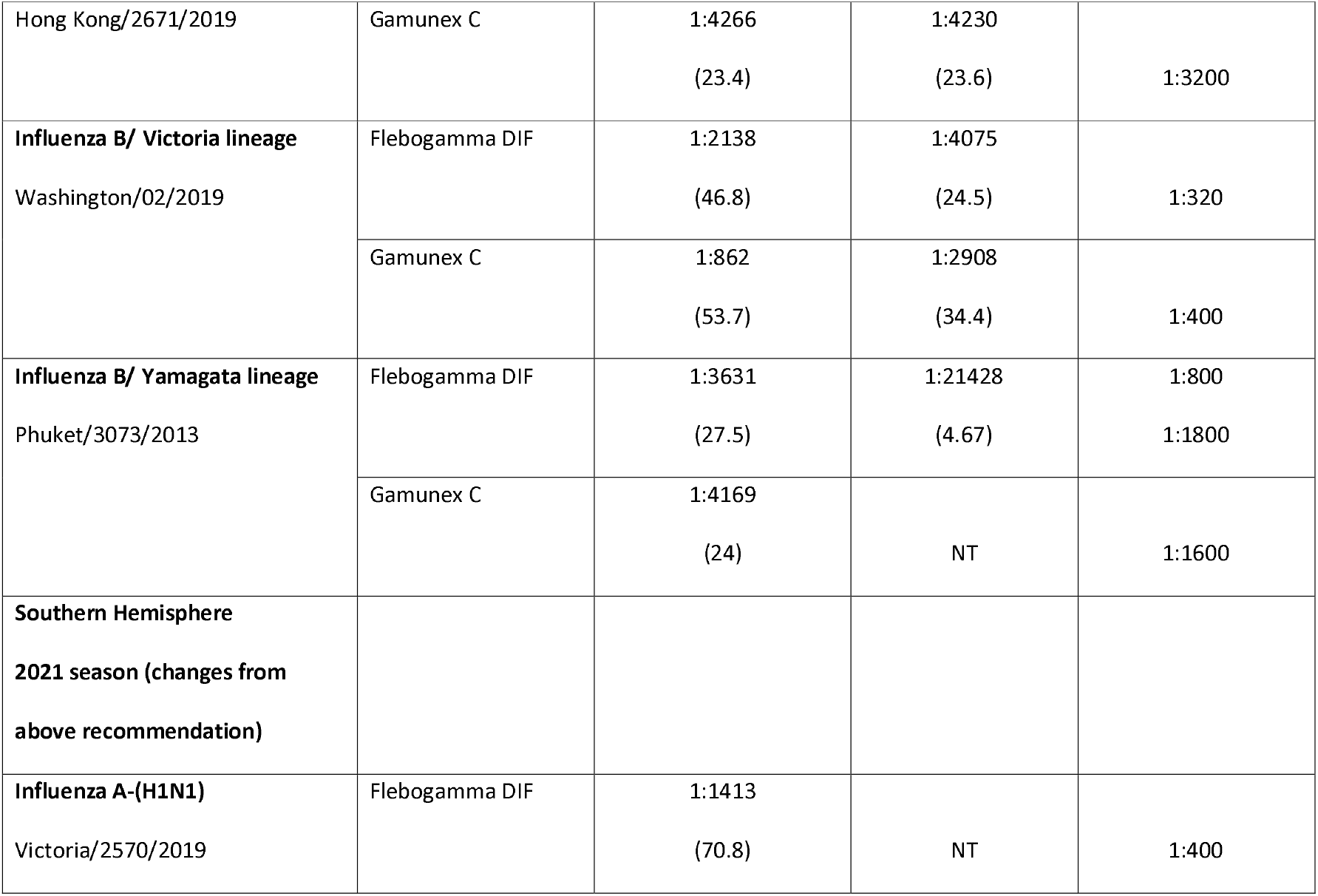
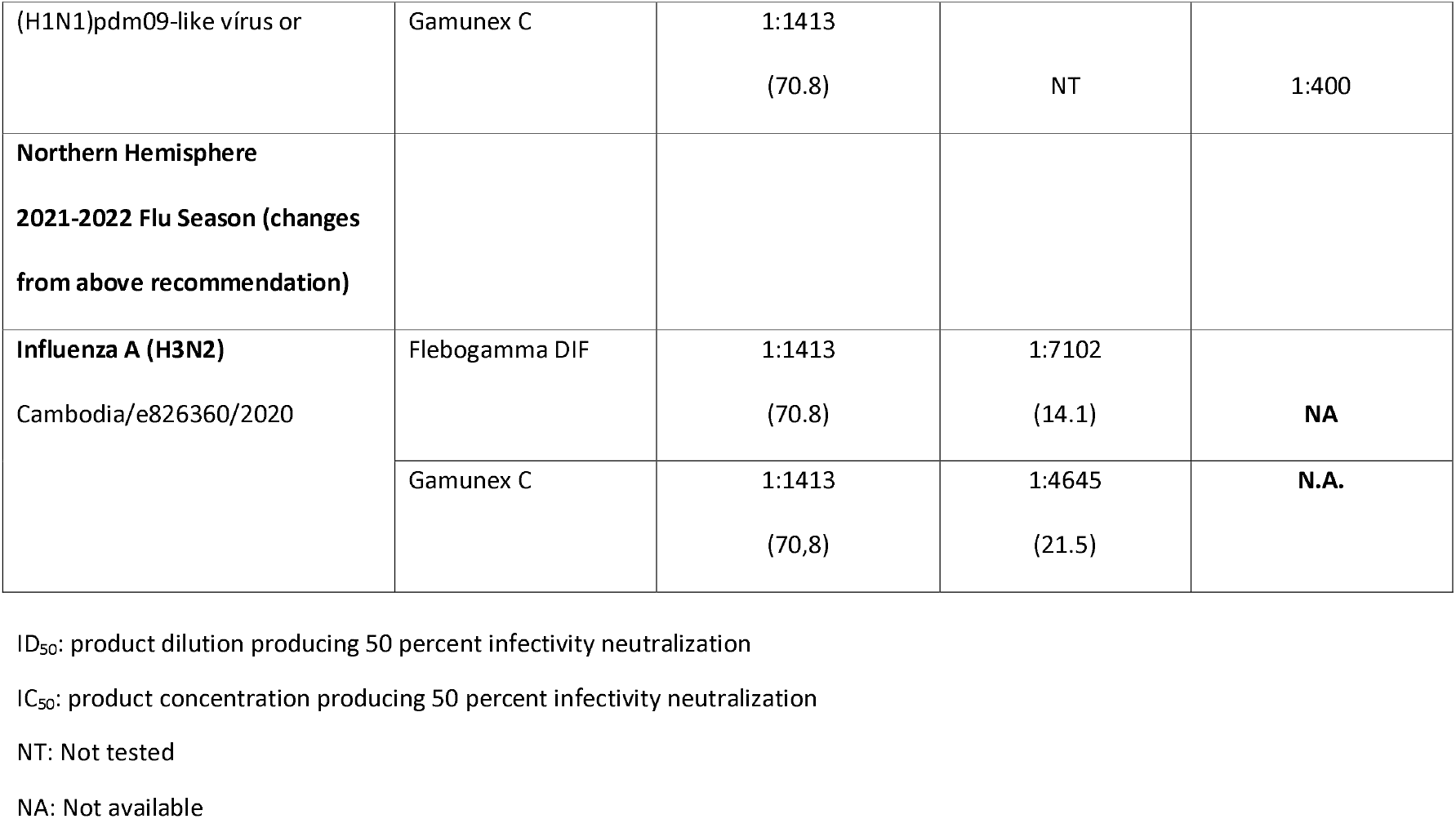
Hemagglutination inhibition and infectivity neutralization titers for seasonal IVIG (2020) against influenza strains recommended by the WHO for 2020-2022 flu seasons in the Northern and Southern Hemispheres.

The concentration-response curves for IVIG (June 2020) neutralization against the WHO-selected influenza strains selected for the Northern Hemisphere 2020-2021 flu season are shown in Figure 1. These curves were used to calculate the IC_50_ values given in Table 1. Neutralization activity (IC_50_) was similar for the selected influenza strains. Neutralization titers were in the range of 1:2399-1:21428 and IC50 values ranged from 4.67-41.7 μg/mL. The higher end of IC_50_ potency was against the influenza B/ Yamagata lineage Phuket/3073/2013 strain: 4.67-24 μg/mL. Potency was similar against three of the other strains (14.7-41.7 μg/mL): influenza A/(H3N2) Hong Kong/2671/2019, influenza B/ Victoria lineage Washington/02/2019 and influenza A-(H1N1) Guangdong-Maonan/SWL1536/2019 strain

**Figure 1:**
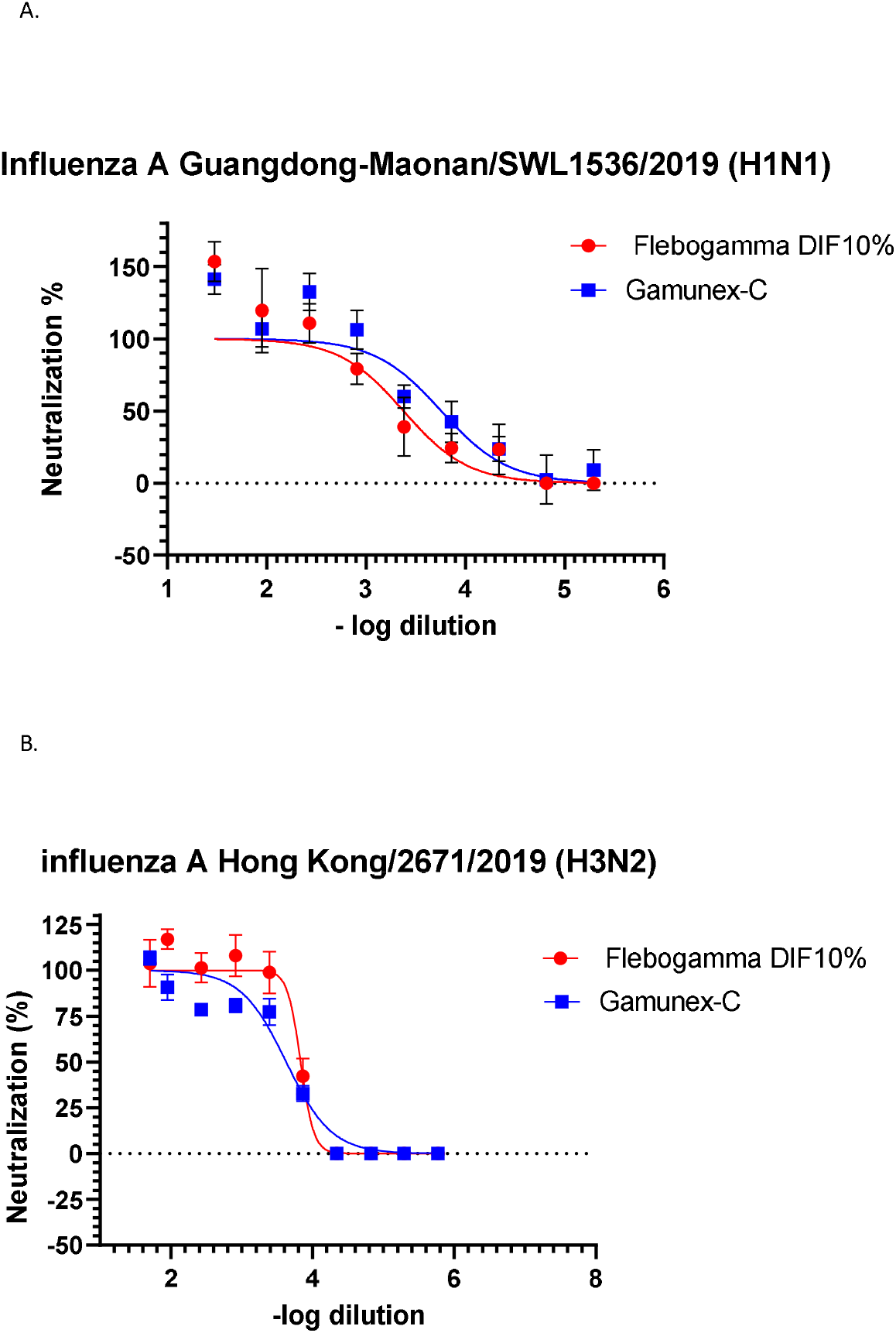
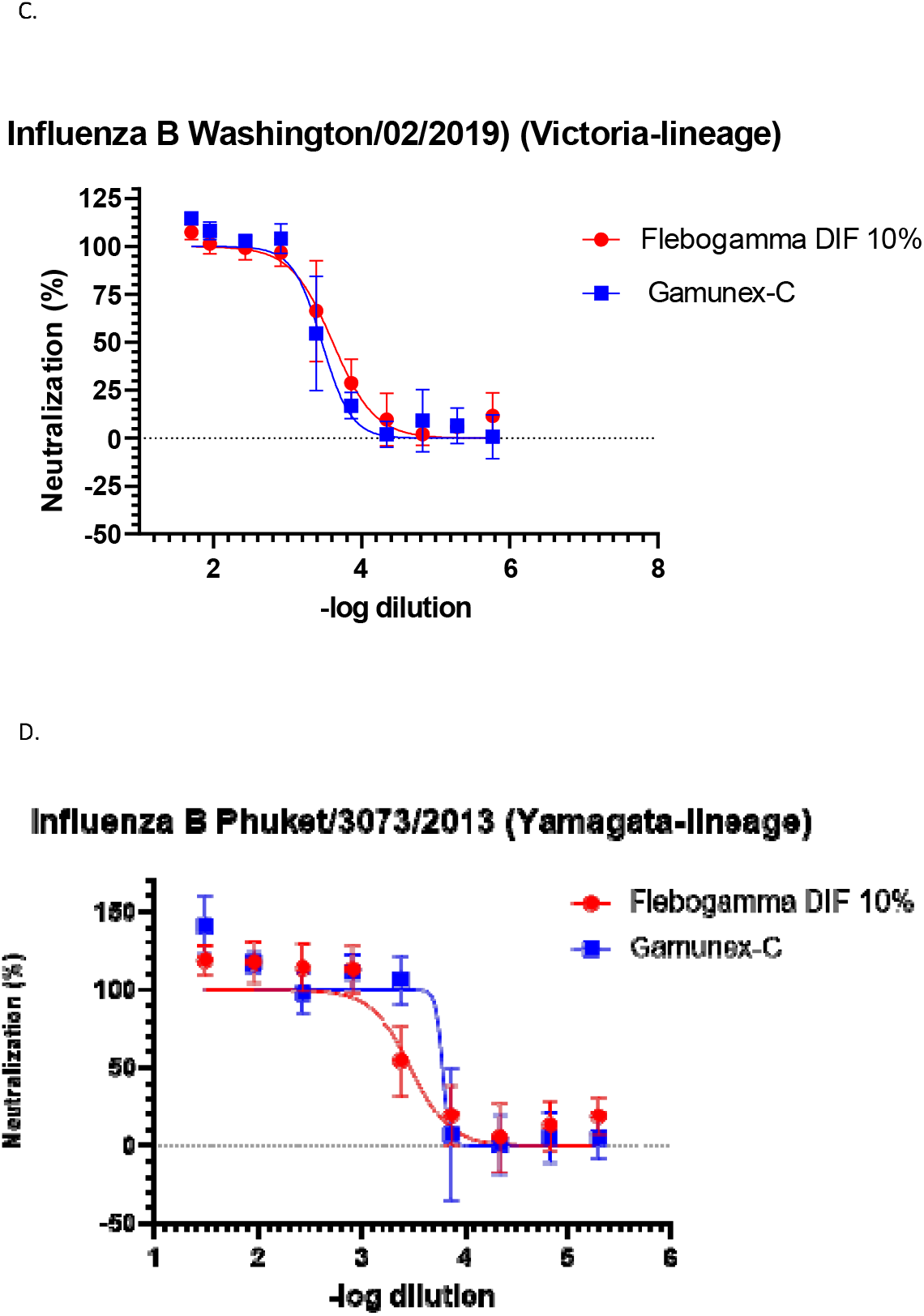
Concentration-neutralization curves for the influenza strains recommended by the WHO for the 2020-21 influenza season ^6^ for neutralization by seasonal IVIG (Flebogamma DIF and Gamunex) manufactured in June 2020. **A.** Effects in the influenza A Guangdong-Maonan/SWL1536/2019 (H1N1)pdm09-like strain. **B.** Effects in the influenza A Hong Kong/2671/2019 (H3N2)-like strain. **C.** Effects in the influenza B Washington/02/2019 (Victoria lineage)-like strain. **D.** Effects in the influenza B Phuket/3073/2013 (Yamagata lineage)-like strain.

Hemagglutination inhibition titers for seasonal IVIG against the selected strains of influenza (Northern Hemisphere 2020-2021) were high: 1:320-1:3200.

The same IVIG product batches (June 2020) showed similar titers against the different H1N1 influenza A virus recommended for the Southern Hemisphere 2021 flu season (Victoria/2570/2019 pdm09-like) and the different H2N3 influenza A virus recommended for the Northern Hemisphere 2021-2022 flu season (Cambodia/e826360/2020).

## Discussion

IVIG and IgG products for other routes of administration (subcutaneous or intramuscular) are produced from pools of human plasma collected from thousands of healthy donors. Our hypothesis was that these plasma pools and the resulting IG products mirror the immunological history of the donor population in terms of their responses to exposure to influenza viruses. These pools reflect a wide variety of donor of different ages and geographic origins each having their own history of influenza exposure. The anti-influenza antibodies in these IG products may arise through a combination of infections and vaccinations over the years in each donor.

Influenza viruses have evolved two different mechanisms (antigenic drift and antigenic shift) to escape natural immunity. These mechanism also lessen the efficacy of influenza vaccines. ^11^ Consequently, surveillance programs have been set up by the WHO to constantly monitor antigenic changes in influenza viruses around the world. Based on this surveillance program, the WHO recommends the viral strains to be targeted by vaccines for the Northern and Southern Hemisphere influenza seasons. As previously mentioned, the strains selected for the quadrivalent vaccine include two A strains and two B strains.

Vaccination strategies are classically aimed at the hemagglutinin surface protein and consequently standard methods to measure anti-influenza antibody titers are directed towards hemagglutination and hemagglutination inhibition. Hemagglutination inhibition detects antibodies against hemagglutinin which are classically considered neutralizing antibodies. Classically, a hemagglutination inhibition titer of 1:40 has been considered protective.^12, 13^ All the hemagglutination titers measured in this study were well above this value. At least theoretically, under normal infusion conditions (200 mg IgG/kg weight), these titers would be sufficient to provide a high enough recipient plasma titer (≥ 1:40) to consider these IVIG solutions to be protective against these influenza viruses

However, there are other proteins on the influenza virion that may be the target of neutralizing antibodies (e.g., neuraminidase). Using cell culture methods to directly measure infectivity neutralization detects not only neutralizing antibodies towards hemagglutinin, but also neutralizing antibodies targeting different influenza surface proteins. Therefore, this methodology would detect neutralizing antibodies to neuraminidase, the target of some antiinfluenza drug therapies.

IVIG solutions manufactured in June 2020 (from plasma collected in the prior 6-months) were tested in the current study for activity against the four influenza strains recommended for the Northern Hemisphere 2020-2021 influenza season,^6^ the strains recommended for the Southern Hemisphere 2021 influenza season ^7^ and the strains recommended for the Northern Hemisphere 2021-2022 season. ^8^ These IVIG solutions were tested using inhibition of hemagglutination and infectivity neutralization methodologies.

As mentioned above, seasonal influenza epidemics vary from year to year but frequently extract a heavy global toll in terms of morbidity and mortality. Vaccination is the mainstay for controlling these influenza epidemics. Additionally, antiviral drugs play an important role in post-exposure prophylaxis and treatment of influenza infections. However, a lack of efficacy and the development of viral resistance can limit the effectiveness of these treatments. Despite these treatments, significant morbidity and mortality can still occur especially in immunocompromised patients. ^14^ Treatment of patients with compromised immune systems (due to hereditary factors, chemotherapy, organ transplantation or other causes) might include prophylaxis as well therapy for active disease.

A recent study found that the breadth of the antibody response from a natural infection was quite different from that produced by vaccination. ^15^ This observation could theoretically increase the diversity of antibodies in IVIG by virtue of inclusion of both infection and vaccination-induced antibodies. The diversity of antibodies along with the well-documented cross-reactivity between antibodies to different influenza virus strains could lead to broad activity for IVIG against influenza infections. The broad-based activity may be reflected in the activity of a single season IVIG against all influenza strains selected for three consecutive influenza seasons.

## Conclusions

In the circumstances of viral resistance and/or a lack of effectiveness of the usual treatments, additional prophylactic or treatment options are needed. The data presented here show that this seasonal IVIG (June 2020) had high hemagglutination inhibition and neutralization titers against the influenza strains selected for inclusion in seasonal flu vaccines. These results raise the potential that IVIG could be used together with the aforementioned treatment strategies to help control seasonal influenza epidemics. Immunocompromised patients, those with comorbidities and the general population could potentially benefit from IVIG treatment. In any case, this possibility deserves and needs more investigation.

## Acknowledgements

The authors acknowledge the expert technical assistance of the personnel of the Immunotherapies Unit, Grifols Global Research & Development Bioscience Industrial Group (Barcelona, Spain), Eduard Sala, Judith Luque and Daniel Fajardo. In addition, Michael K. James, Ph.D.is acknowledged for medical writing and Jordi Bozzo, Ph.D., CMPP for editorial assistance.

## Notes

**Conflicts of interest:** JMD, DC, CR and RG are employees of Grifols.

### Competing Interest Statement

José María Díez, Daniel Casals, Carolina Romero and Rodrigo Gajardo are employees of Grifols.

