## Supplemental Material for "Seasonal IVIG contains high infectivity neutralizing and hemagglutination inhibition activity against seasonal flu virus strains selected for vaccines (2020-22)"

### **Supplementary Material**

#### **Microneutralization methods**

The microneutralization methodology used in this study was previously described^1^. Viral suspensions 100 IU/125 µL (800 IU/mL) were prepared in Dulbecco’s modified Eagle Medium (DMEM) with 2% fetal bovine serum (FBS). Dilutions of IVIG were prepared in the same medium (DMEM with 2% FBS). The viral suspension (125 µL) was added to the wells of a 96-well plate except for the wells designated as negative controls and toxicity controls. These control wells received 125 µL of the dilution medium (DMEM 2% FBS). Nine dilutions of the test IVIG solutions were added to test wells (eight replicates). The most concentrated IVIG dilution was added to the eight toxicity control wells.

After incubation for 90 min at 37 ± 2°C, a 200 µL aliquot of the suspensions described above was transferred to a 96-well plate containing Madin-Darby bovine kidney cells (MDBK) grown to near confluence. After addition of the suspensions, a second incubation period was carried out (37 ± 2°C, 8% CO_2_) until a generalized cytopathic effect was seen in all the positive control wells.

The results after the second incubation were read using a luminometer (Tecan Infinite^®^ 200 Pro, Model M Plex, Männedorf, Switzerland) using the Viral ToxGlo^TM^ reagent (Promega, Madison, WI, USA) for the cytotoxicity effect or by CPE observed under an inverted microscope (Zeiss, Axiovert ; 40, Achroplan 10X/0.25 Ph1 objective. Zeiss, Oberlochen, Germany) followed with TCID_50_ titer calculation.

**Reference**:

1.	WHO Global Influenza Surveillance Network: Manual for the laboratory diagnosis and biological surveillance of influenza. World Health Organization; 2011.
